# Contrasting response of two *Lotus corniculatus* L. accessions to combined waterlogging-saline stress

**DOI:** 10.1101/2020.04.28.066621

**Authors:** C.J. Antonelli, P.I. Calzadilla, M.P. Campestre, F.J. Escaray, O.A. Ruiz

**Author notes:** These authors contributed equally to this work (C.J. Antonelli, P.I. Calzadilla). <u>Corresponding author:</u> Oscar Adolfo Ruiz;<u>; Tel:</u> (0054) - 2241 430323. Instituto de Fisiología Vegetal (INFIVE), Universidad Nacional de La Plata (UNLP) y Consejo Nacional de Investigaciones Científicas y Técnicas (CONICET), Diagonal 113 No. 495, 1900 La Plata, Argentina. Department of Earth and Environmental Sciences, Faculty of Science and Engineering, University of Manchester, Manchester M13 9PT, UK.

## Abstract

Waterlogging and salinity impair crops growth and productivity worldwide, being their combined effects larger than the additive effects of both stresses separately. Recently, a new *Lotus corniculatus* accession has been collected from a coastal area with a high frequency of waterlogging-saline stress events. This population is diploid and has potential to increase nutritional values of *Lotus* cultivars used as forages. Due to its environmental niche, we hypothesize that this accession would show a better adaptation to combined waterlogging-saline stress compared to another commonly used tetraploid *L. corniculatus* (cv. San Gabriel). Shoot and root growth under waterlogging, salinity and combined waterlogging-saline treatments were addressed, together with chlorophyll fluorescence and gas exchange measurements. Results showed that salinity and waterlogging effects were more severe for the tetraploid accession, being the differences larger under the combined stress condition. In addition, Na^+^, Cl^−^ and K^+^ concentrations were measured in young and old leaves, and in roots. A larger accumulation of Na^+^ and Cl^−^ was observed under both saline and combined stress treatments for the tetraploid *L. corniculatus*, for which ion toxicity effects were evident. The expression of the *NHX1* and *CLC* genes, coding for Na^+^ and Cl^−^ transporters respectively, was only increased in response to combined stress in the diploid *L. corniculatus* plants, suggesting that ion compartmentalization mechanisms were induced in this accession. As a conclusion, the recent characterized *L. corniculatus* might be used for the introduction of new tolerant traits to combined stresses, in other *Lotus* species used as forage.

## 1. Introduction

Human agriculture is facing one of its most important challenges due to an increase in food demand and the more severe consequences of global climate change (Thornton et al., 2014). Under these circumstances, the decrease in biological diversity and widespread monoculture add more uncertainties to the problem (Nunez et al., 2019). In this context, discovering new germplasm adapted to restrictive soils or environments will be of fundamental importance, allowing the introduction of new valuables traits to the existing crop species.

Within climate change consequences, global warming alters the precipitations regime, increasing the frequency of flooding and drought events worldwide (Hirabayashi et al., 2013). This phenomenon, together with the sea-level rise in coastal areas (Carter et al., 2006; Martin et al., 2011) and the expansion of high sodicity soils (Ghassemi et al., 1995; Lavado and Taboada, 1987), also increase the frequency of combined waterlogging-saline stress events (Bennett et al., 2009).

The effects of waterlogging and salinity stress have been extensively studied in different plant species. Waterlogging limits oxygen diffusion in the rhizosphere (Armstrong and Drew, 2002; Ponnamperuma, 1984), compromises mitochondrial respiration in roots (Gupta et al., 2009; Zabalza et al., 2009) and leads to an ATP production failure for energy demanding processes (Bailey-Serres and Voesenek, 2008; Geigenberger, 2003). Thus, plant responses to waterlogging involve anatomical and morphological changes, such as aerenchyma formation and development of adventitious roots, which aim to increase root oxygenation (Colmer and Voesenek, 2009; McDonald et al., 2002). In particular, in the *Lotus* genus, aerenchyma formation was shown to correlate with flooding tolerance in several species (Antonelli et al., 2019; revised by Striker and Colmer, 2017).

Regarding salinity, salt causes osmotic stress and ionic toxicity in most crop plants (Blumwald, 2000; Munns, 2002). Firstly, the osmotic stress occurs as a consequence of the decrease of water availability in roots due to the increase in ion concentration in the soil. Ion toxicity take place in a second phase, and is caused when Na^+^ and/or Cl^−^ are accumulated in leaves, disrupting protein structure, organelles and affecting cell metabolism (reviewed by Munns and Tester, 2008). While the osmotic stress effect is immediate, the effect due to ionic toxicity is observed after longer exposure time (days or weeks) (Munns, 2002). In addition, salinity may cause nutrient deficiencies or imbalances, due to the competition of Na^+^ and Cl^−^ with nutrients such as K^+^, Ca^2+^and NO_3_^−^ (Hu and Schmidhalter, 2005).

Plants present different strategies to respond to salinity, such as ion exclusion or compartmentalization mechanisms (Munns and Tester, 2008; Teakle and Tyerman, 2010). In the case of the ion exclusion, these mechanisms can be achieved by exclusion transporters, which avoid the entrance of toxic ions into root cells. By contrast, in the compartmentalization mechanisms, ions are accumulated in sub-compartments within the plant cells, avoiding toxic levels to be reached in the cytoplasm and maintaining ionic homeostasis (Munns and Tester, 2008; Teakle and Tyerman, 2010). In both cases, the active transport of ions, with ATP consumption, is required (Munns and Tester, 2008; Teakle and Tyerman, 2010).

When waterlogging occurs together with salinity, the combined effects are larger than the additive effects of both stresses separately (Barrett-Lennard, 2003; Teakle et al., 2010). For instance, the energy deficit caused by waterlogging-induced hypoxia will have a direct impact on the Na^+^ transporters (Byrt et al., 2007; James et al., 2006), the Na^+^/H^+^ antiporters in the plasma membrane (Martínez-Atienza et al., 2007), and/or the Na^+^/H^+^ antiporters of the tonoplast (like NHX1; reviewed by Pardo et al., 2006; Xue et al., 2004). In this sense, it was reported that combined stress largely increases the concentrations of Na^+^ and Cl^−^ in plant shoots, as compared with salinity stress alone (Barrett-Lennard, 2003; Barrett-Lennard and Shabala, 2013).

Between the regions affected by combined waterlogging-saline stress events, it is possible to mention the Flooding Pampa (Argentina). This area comprises approximately 9 million hectares, being one of the most important cattle rearing area in South America (Soriano et al., 1991). Its soil is characterized by a poor nutrient availability, high clay content, salinity and alkalinity. Its frequent exposure to flooding periods, makes the Flooding Pampa a very restrictive environment for crops growth and forage production. As a consequence, the main forage source for cattle bearing consists of natural grasses which are reduced in protein content (Perelman et al., 2001).

Different strategies to improve forage quality traits are carried out in regions like the Flooding Pampa, using plant species adapted to constrain conditions. Within them, the use of species of the *Lotus* genus is in the spotlight due to its high plasticity and nutritional value, being relevant forage alternative in South America, Australia and Europe (Blumenthal and McGraw, 1999; Escaray et al., 2012). In particular, *L. corniculatus* has been extensively used due to its moderate level of proanthocyanidins (PA), which contributes positively to nutritional quality of ruminant diet, increasing protein fraction assimilation, avoiding cattle bloat and reducing intestinal parasites (Foo et al., 1996; McNabb et al., 1996; Min et al., 2003). Nevertheless, the extended use of commercial cultivars of *L. corniculatus* (all tetraploids) has failed due to its edaphic requirements and its susceptibility to different stress conditions such as flooding or salinity (Antonelli et al., 2019; Escaray et al., 2019). By contrast, *L. tenuis* developed a relatively greater tolerance to waterlogging and salinity conditions (revised by Striker and Colmer, 2017), becoming naturalized in the Flooding Pampa. This species presents forage quality comparable to that of other forage legumes (such as *Medicago spp*. or *Trifolium spp*.). However, the low productivity of *L. tenuis* and its PA absence at foliar level, make of *L. corniculatus* a better forage species (revised by Escaray et al., 2012).

Recently, a new *L. corniculatus* accession has been collected from an alkaline-salty area, in the Valencia Albufera in Spain (Escaray et al., 2014). This population is diploid and has been used to improve *Lotus* cultivars through inter-specific hybridization. Although this new germplasm was tested under waterlogging stress (Antonelli et al., 2019) and salinity (Escaray et al., 2019), its stress response under combined waterlogging-saline stress was not yet evaluated. In the present study, we aimed to compare the response to waterlogging, salinity and combined waterlogging-saline stress of the diploid accession of *L. corniculatus* (described by Escaray et al., 2014) and one of the most common *L. corniculatus* cultivars used as forage production in South America (L. *corniculatus* cv. San Gabriel). Due to its environmental niche, we hypothesize that the diploid accession would show a better adaptation to combined waterlogging-saline stress than the commercial *L. corniculatus* cultivar.

## 2. Materials and Methods

### 2.1 Plant material

The *L. corniculatus* diploid accession (LcD) corresponds to a wild population collected at the Devesa de El Saler, Valencia (Spain). The taxonomical identity of this population as belonging to *L. corniculatus* was established and confirmed by different authors (Ballester Ramírez de Arellano, 2015; Escaray et al., 2014). *L. corniculatus* commercial cv. San Gabriel (LcT) is a germplasm obtained by INIA (Instituto de Investigación Agropecuaria de Uruguay, Uruguay).

### 2.2 Experimental design and growth conditions

A completely randomized design was performed and the two mentioned plant materials were evaluated under four treatments: 1-control treatment, plants were irrigated with nutrient solution and free drainage (Ctrl); 2-waterlogging with nutrient solution, keeping the water column 3 cm above the pot substrate surface without drainage (WL); 3-salinity, nutrient solution with 150 mM of NaCl and free drainage (NaCl); and 4-combined waterlogging-saline, a combination of the two last treatments (no drainage) (WL+NaCl).

Experiments were initiated from seeds, which were scarified with concentrated sulfuric acid (98%) during 3 min, washed ten times with sterile distilled water and sown in Petri dishes containing water-agar (0.8%). Plates were incubated for 7 days in a growth chamber, with a 16/8 h photoperiod at 24/21 ± 2°C (day/night) and 55/65 ± 5% relative humidity. Light intensity (250 μmol photons m^-2^ s^-1^) was provided by Grolux fluorescent lamps (F 40W). Seedlings at the full expanded cotyledon stage were transferred to 300 cm^3^ pots, containing a mixture of washed sand-perlite (1:1 V/V), and irrigated as explained above. An individual plant per pot was considered an experimental unit. After 21 days of growing, when plants showed five fully-developed leaves, the stress treatments were initiated. Irrigation was performed throughout the experiment with a modified 0.5 x Hoagland’s nutrient solution (Hoagland and Arnon, 1950) containing 3 mM KNO_3_; 2 mM Ca(NO_3_)_2_.4H_2_O; 1 mM MgSO4.7H2O; 0.5 mM NH_4_H_2_PO_4_; 50 μM NaFeO_8_EDTA.2H_2_O; 4.5 μM MnCl_2_, 23 μM H3BO3, 0.16 μM CuSO_4_.5H_2_O, 0.09 μM ZnSO_4_.7H_2_O, and 0.06 μM Na_2_MoO_4_.2H_2_O.

### 2.3 Stress treatments

In order to avoid any osmotic shock in saline and combined waterlogging-saline treatments, plants were treated with increasing concentration of NaCl (starting from 25 mM and reaching a final concentration of 150 mM) during 12 days (acclimation period). Overall, plants were subjected to NaCl during 33 days. In all pots subjected to waterlogging, nutrient solution containing 0.1% (w/v) agar was bubbled with N_2_ gas before irrigation. This procedure lowers the dissolved O_2_ levels to less than approximately 10% of air-saturated solution (Gibbs and Greenway, 2003), whereas the dilute agar prevents convective movements in the solution (Wiengweera et al., 1997). There were five pots per treatment (n = 5), and the experiments were repeated at least once. Data shown corresponds to the most representative experiment.

### 2.4 Plant growth measurement

At harvest, plant tissues were divided in young leaves (upper 2 fully expanded leaves), old leaves, stems and roots, and their dry matter was determined after drying at 60°C, until constant weight. By adding the weight of different tissues, total dry weight and the shoot:root ratio were calculated.

### 2.5 Photosynthesis measurements

One day before harvest, net photosynthetic rate at light saturation (Asat) was measured on the second apical fully expanded leaf (1500 μmol photons m^-2^ s^-1^ illumination, LED light), using a portable photosynthesis system (TPS-2 Portable Photosynthesis System, MA, USA). Then, net photosynthetic rate was relativized to leaf area. For this purpose, leaves were scanned and their area estimated using an image analyser program (Image Pro Plus 4.5). Non-invasive OJIP test (Strasser and Srivastava, 1995) was also performed on the second fully expanded leaf using a Pocket PEA Chlorophyll Fluorimeter (Hansatech Instruments, UK). Leaves were dark adapted for 20 min before analysis and then exposed for 3 s to light at an intensity of 3500 μmol m^-2^ s^-1^. Data were processed by PEA Plus software (Hansatech Instruments, UK) and Windows Excel (Microsoft, WA, USA). The maximum quantum yield of primary photochemistry (Fv/Fm) was calculated.

### 2.6 Analytical determinations

An aliquot of 10 mg of dried material was used to estimate the concentration of Na^+^ and K^+^ by standard flame photometry (Proehl and Nelson, 1950). Chloride was determined by a thiocyanate-Hg-based colorimetric reaction (Iwasaki et al., 1956). For this, 12.5 mg of powdered dry plant material was extracted in 0.5 mL of a solution containing H_2_O_2_ (30%):concentrated HNO_3_:isoamyl alcohol:H_2_O at 1:1:0.08:7.9 (V/V). The extraction was incubated at room temperature for 15 min, diluted to 5 mL with Milli-Q. water and vigorously agitated in a Vortex. Then, 1.5 mL of the extraction mixture was centrifuged (10,000 rpm, 5 min) and the supernatant transferred to another tube. The colorimetric reaction solution contained polyethylene glycol dodecyl ether–water (Brij 35®, 4%):mercuric thiocyanate (4.17 g/L methanol):(NO3)3Fe (202 g/L Milli-Q. water plus 21 mL concentrated HNO_3_):Milli-Q water at 0.05:15:15:70 (V/V). For treatments not involving NaCl, one millilitre of colorimetric reaction was added to 320 μL of the sample supernatant. For treatments that include NaCl, 50 μL of the supernatant were previously diluted with extraction solution to 320 μL. Sample absorbance was determined at 450 nm with a spectrophotometer (Hitachi U-1100), and interpolated into a KCl calibration curve (0, 5, 10, 15, 20 ppm).

### 2.7 RNA isolation and cDNA synthesis

Total RNA was extracted from frozen apical leaves using a Plant Spectrum Total RNA Kit (Sigma), according to the manufacturer’s instructions, and treated with DNase (TURBO DNA-free™ Kit, Ambion). The quality and quantity of RNA were verified by agarose gel electrophoresis and spectrophotometric analysis. The absence of DNA from the RNA samples was tested by the null PCR amplification of the universal rDNA primer pair ITS1/ITS4, as described in Paolocci et al (2006). Then, cDNA from *L. corniculatus* plants was synthesized from 3 μg of total RNA using a Moloney Murine Leukemia Virus Reverse Transcriptase (MMLV-RT) (Promega, WI, USA) and 100 pmol of random hexamers (Pharmacia Biotech), according to supplier’s instructions.

### 2.8 Primer Design

The cDNA sequences related to the *CLC* genes from *L. Japonicus* and other model species were downloaded from GenBank. Using own transcriptomic information from *L. corniculatus* species (manuscript in preparation), an *in silico* analysis was performed to obtain a *CLC* homologous sequence for *L. corniculatus*. Deduced *CLC* sequences from *L. corniculatus* and *L. Japonicus* were aligned with the BioEdit program (Hall, 1999). The most conserved nucleotide sequence between species was used to design the corresponding primers for qRT-PCR, by the software Primer3Plus (http://www.bioinformatics.nl/cgi-bin/primer3plus/primer3plus.cgi). Additionally, a primer pair reported by Teakle et al. (2010) was used to evaluate the relative expression of the *NHX1* gene. In all cases, the *EF-1α* gene was used as housekeeping (Escaray et al., 2014).

The primer pairs were initially checked for their specificity and amplification efficiency in both *L. corniculatus* accessions. Only primer pairs that produced the expected amplicon and showed similar PCR efficiency were used in the present study. Primers used for qRT-PCR analysis are listed in Supplementary Table 1.

### 2.9 Quantitative RT-PCR

An aliquot of 5 μL of 1:8 diluted cDNA was used in the qRT-PCR reactions, made using 15 μL of the FastStart Universal SYBR-Green Master Mix (Rox, Roche) and 2.5 pmol of each primer, according to the supplier’s instructions. Three biological replicates were performed per sample and gene. Cycling parameters were two initial steps of 50°C for 2 min and 95°C for 2 min, a two-step cycle of 95°C for 15 s and 60°C for 1 min repeated 50 times, and a final step of 10 min at 60°C. This was followed by the dissociation protocol. Amplifications were performed on Mx3005P qPCR System apparatus with the help of the MxPro qPCR Software 4.0 (Stratagene, La Jolla, CA, U.S.A.). For each transcript, the average threshold cycle (Ct) was determined. The gene quantification method was based on the relative expression of the target gene versus the reference *EF-1α* gene, according to Paolocci et al. (2007).

### 2.10 Statistical analysis

The experimental results were analyzed using one-way ANOVA for each of the two accessions independently of each other, with four treatments (Ctrl, WL, NaCl and WL+NaCl), followed by Duncan’s test (p < 0.05). Previously, the assumptions of variance homogeneity and normality were tested for all variables with Levene’s and Shapiro-Wilk, respectively. In the case of the analysis of the maximum quantum yield of PSII (Fv/Fm), a t-test was performed. In some cases, comparison between the accessions was also performed using a t-test analysis, for the determined growth variables and ion accumulation measurements. Statistical analysis of gene relative expression was performed based on the pairwise fixed reallocation randomization test (p < 0.05) (Pfaffl et al., 2002). In all cases the Infostat software tool was used (Di Rienzo et al., 2010).

## 3. Results

### 3.1 Effect of salt, waterlogging and combined waterlogging-saline stress on the growth of the two *L corniculatus* accessions

The stress treatments imposed visually affected both *L. corniculatus* accessions after 20 days (12 d of salt acclimation plus 8 d of full treatments) (Figure 1). Effects of saline and waterlogging-saline treatments were more evident for LcT plants, for which a smaller number of stems and leaves was observed when compared to control. Similar symptoms were also observed for this accession under waterlogging, although this effect was less strong when compared to the other stress treatments. Regarding LcD, the stress treatments did not reduce the growth of the plants during the first week of the experiment.

**Figure 1.**
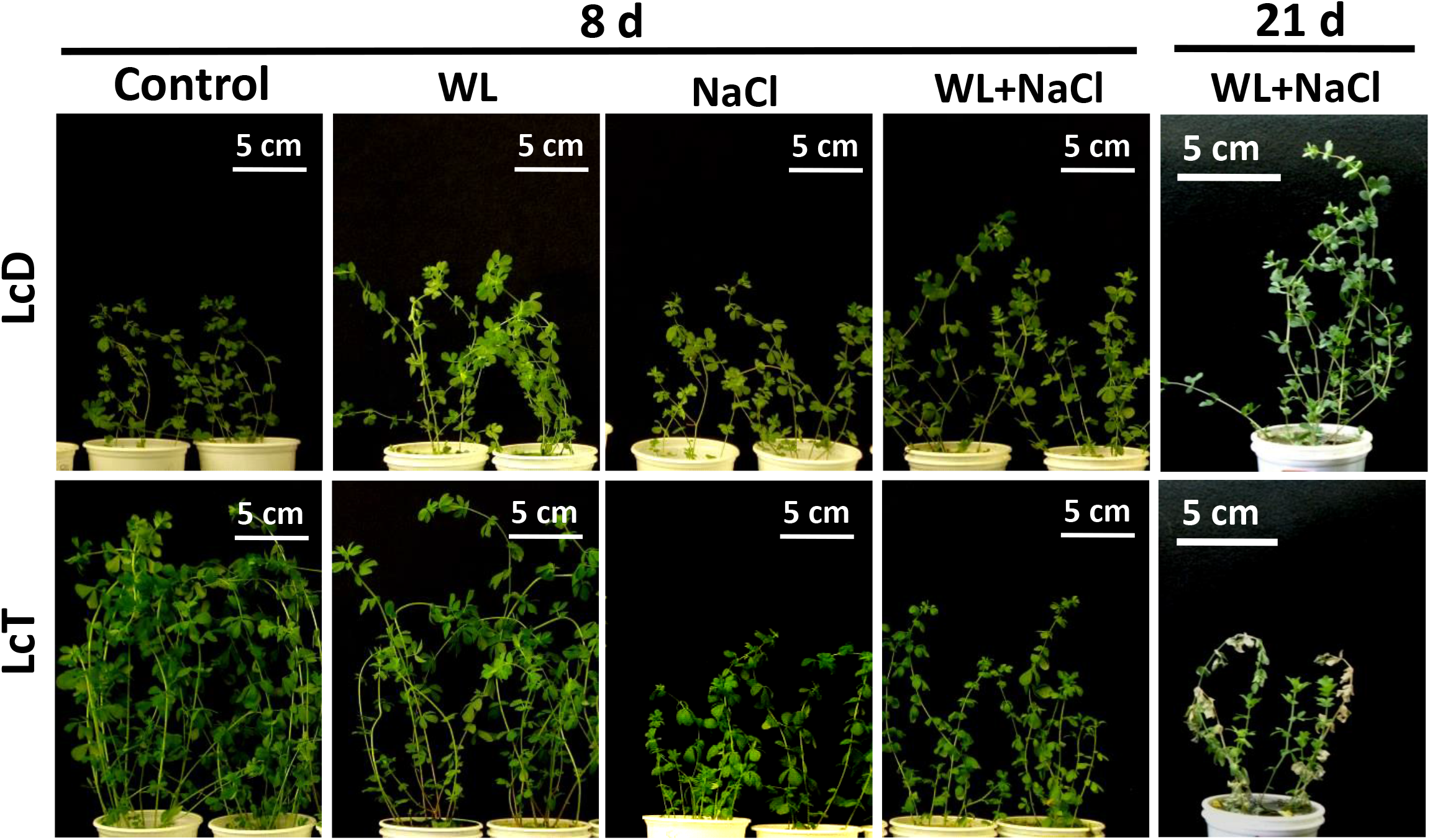
Phenotypic characteristics of *L. corniculatus* accessions subjected to the different stress treatments. Plants were subjected to the different treatments for 21 days, after the salinity acclimation period imposed for those treatments involving salt stress. Pictures were taken after 8 and 21 d (harvest). For the control and waterlogging (WL) treatments, plants were irrigated periodically with Hoagland solution 0.5 x with or without free drainage, respectively. In the salt treatment (NaCl) and combined stress treatment (WL+NaCl), plants were irrigated with Hoagland solution 0.5 x supplemented with 150 mM of NaCl with or without drainage, respectively. When waterlogging stress was imposed, nutritive solution was previously bubbled with N_2_ (g) to reduce O_2_ to hypoxic levels. LcD, diploid *L. corniculatus* accession; LcT, tetraploid *L. corniculatus* accession.

After 33 d since the beginning of the experiment (12 d of salt acclimation plus 21 d of full treatments; harvest date), the phenotype of the plants subjected to the combined waterlogging-saline stress was markedly different between the studied accessions (Figure 1), being LcT more severely affected than LcD. While dead leaves and chlorosis were clearly observed for LcT, none of these symptoms were registered for LcD. The phenotype of the others treatments followed the same trend observed for day 8 (not shown).

The effect of the different stress treatments on the growth of both *L. corniculatus* accessions can also be observed through their accumulated shoots and roots dry mass (Figure 2). Although the variability between biological replicates was larger for LcD than for LcT, no significant differences in shoots and roots dry mass accumulation were observed for LcD among stress treatments and its control (Figure 2A and C). By contrast, the stress treatments significantly reduced the shoots and roots dry mass of LcT. This effect was more pronounced in the combined stress treatment, when compared with the waterlogging and salinity conditions imposed independently. The dry mass reduction for LcT between the combined stress treatment and the control was of 75 % and 60 % for shoots and roots, respectively. It is worth mentioning that the total dry mass accumulation was higher for LcT than for LcD (t-test, *p* < 0.05), in control, waterlogging and saline treatments (Figure 2B). However, under the combined stress treatment, both accessions accumulated similar amount of total dry mass (t-test, *p* > 0.05)

**Figure 2.**
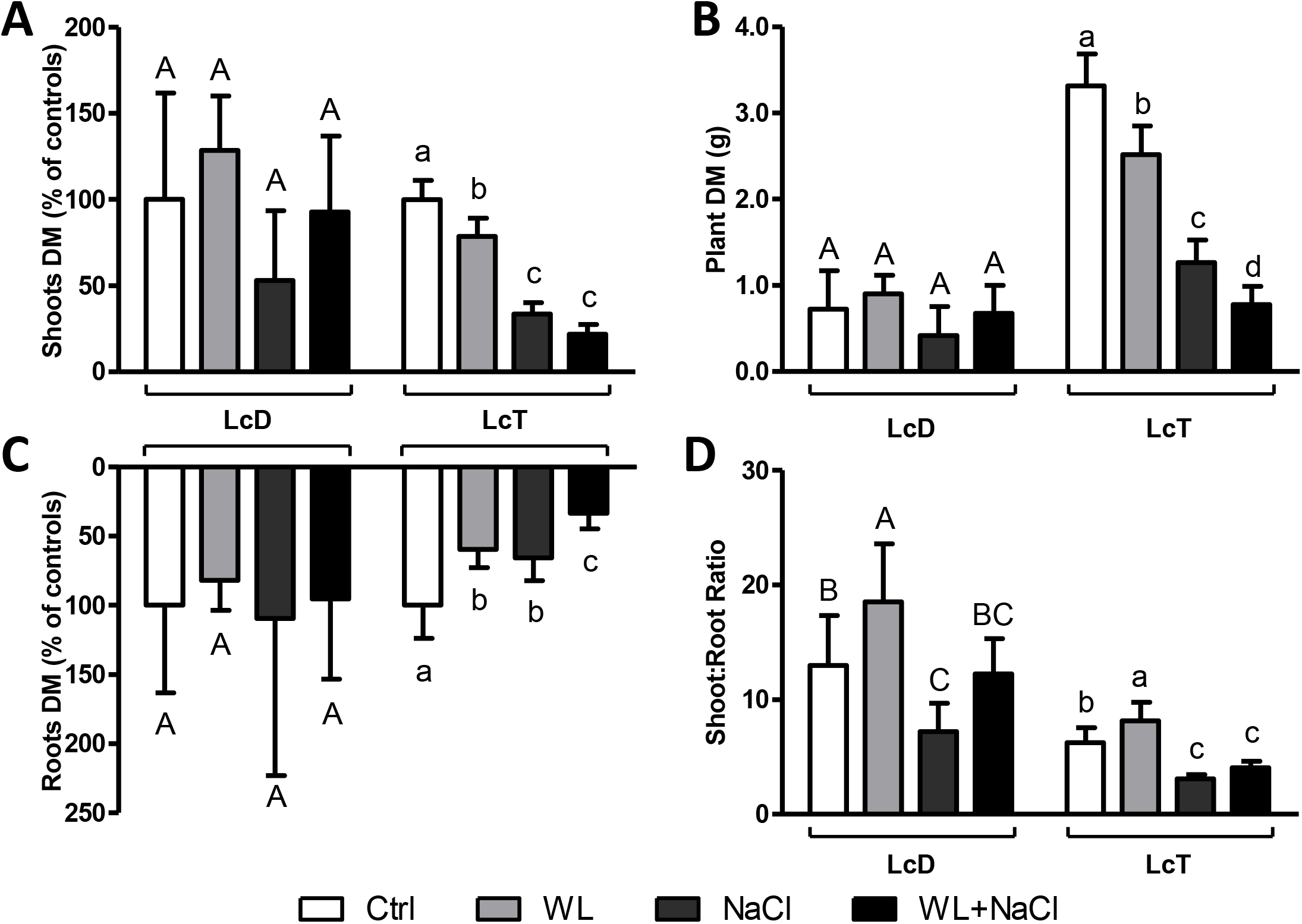
Dry mass accumulation of *L. corniculatus* accessions subjected to the different stress treatments. The measured shoots (A) and roots (C) dry mass are shown as percentage of the control dry mass in each case. (B) Total dry mass (g) accumulation at the end of the experiment. (D) Shoot:Root ratio was calculated from the dry mass accumulation of shoots and roots, respectively. Means (n = 5 ± SD) without common letters differ significantly within each of the accessions (one-way ANOVA; Duncan, *p* < 0.05). White bars, control conditions (Ctrl); light gray bars, waterlogging stress (WL); dark gray bars, saline stress (NaCl); black bars, combined stress (WL+NaCl).

The shoot:root ratio was calculated in the different treatments, from the dry mass accumulated for each accession (Figure 2D and Supplementary Figure 1). An increased in this parameter was observed for both LcD and LcT as a response to waterlogging, when compared to controls. By contrast, a decreased in shoot:root ratio was determined under salinity for both accessions. Regarding the combined stress treatment, a significant decreased compared to controls, was only observed for LcT.

### 3.2 Photosynthetic response of the LcT and LcD accessions to the different stress treatments

Net photosynthetic rate at saturating irradiance (Asat) and the maximum quantum yield of PSII (Fv/Fm) were measured in all plants one day before harvest date (Figure 3). The stomatai conductance (gs) was also measured in each case (Supplementary Figure 2). Asat values were only reduced for LcD under salt stress, while for LcT a decreased was observed in all the stress treatments. In LcT, the stronger effect was observed under the salt and the combined stress. Differences observed in gs were comparable to the ones showed by Asat. The larger reduction of gs was observed under the saline stress for both LcD and LcT. However, no effect was observed in gs in neither of the plant accessions under the waterlogging stress, when compared to controls. In the case of the combined stress treatment, the gs value decreased strongly for LcT, but not for LcD.

**Figure 3.**
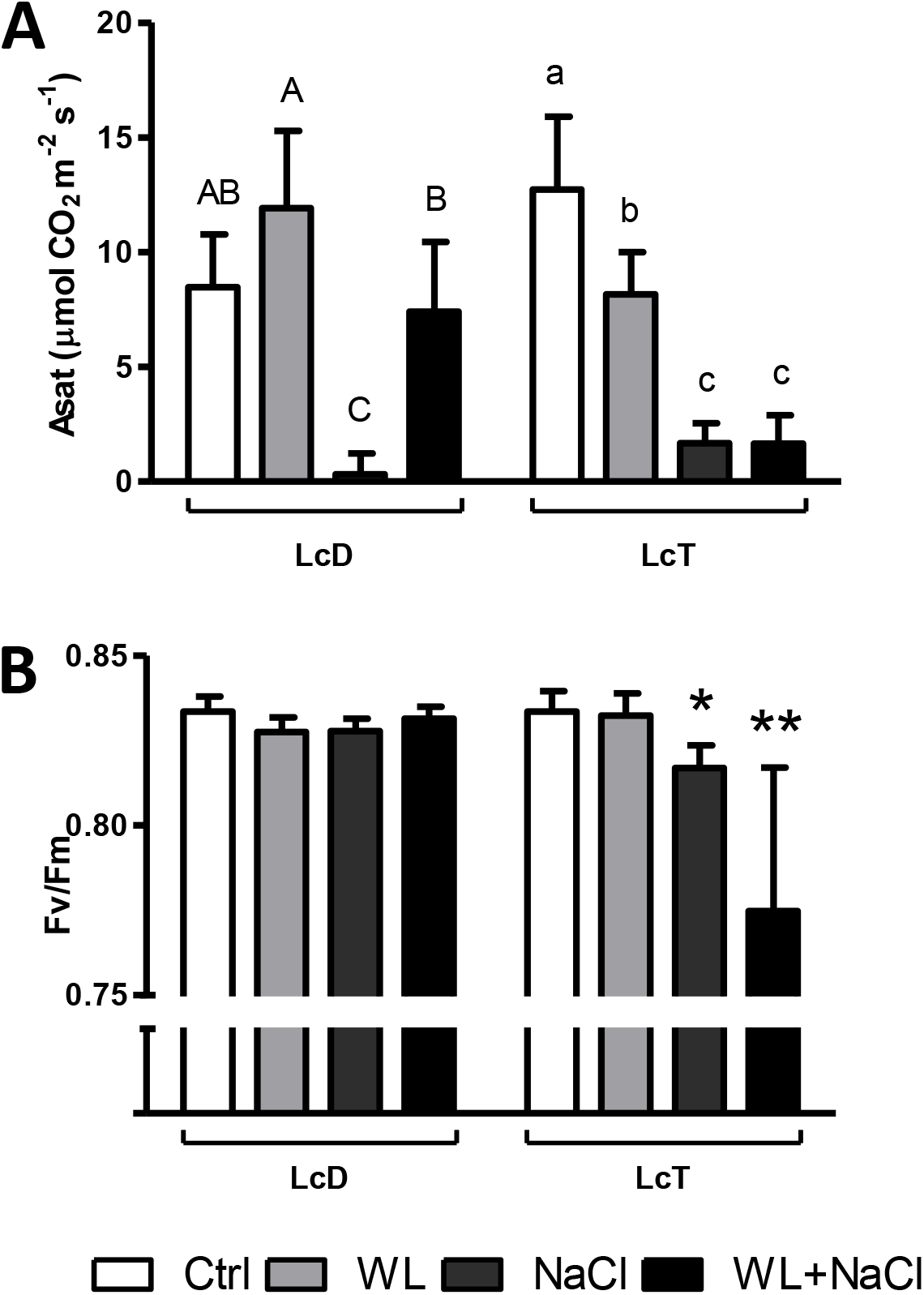
Photosynthetic parameters of *L. corniculatus* accessions under the different stress treatments. Net photosynthetic rate at saturating irradiance (Asat) (A) and the maximum quantum yield of PSII (Fv/Fm) (B) were measured for five independent biological replicates (n = 5 ± SD). Means without common letters differ significantly within each of the accessions (one-way ANOVA; Duncan, *p* < 0.05) (A). Asterisks show significant differences of a stress treatment against the control treatment (Student t-test, * *p* < 0.05; ** *p* < 0.01) (B). White bars, control conditions (Ctrl); light gray bars, waterlogging stress (WL); dark gray bars, saline stress (NaCl); black bars, combined stress (WL+NaCl).

Regarding the Fv/Fm measurements, no differences were observed for LcD between the different treatments; meanwhile, the maximum yield of PSII was significantly reduced for the salt and combined stress condition in LcT, compared to its control (Figure 3B). Reduction of this parameter was more pronounced under the combined stress condition, reaching values lower than 0.8 (Figure 3B).

### 3.3 Ions accumulation in different tissues of LcT and LcD subjected to stress conditions

The concentration of Cl^−^, Na^+^ and K^+^ were measured in apical and basal leaves and in roots of LcT and LcD, at the end of the experiment (harvest date) (Figure 4A-I). The Na^+^/K^+^ was also calculated as a salinity tolerance index, due to the fact that Na^+^ can interfere with K^+^ homeostasis, affecting several metabolic processes (Figure 4J-L) (Assaha et al., 2017; Shabala and Pottosin, 2014). Cl^−^ and Na^+^ concentrations were not altered in the waterlogging treatment when compared with controls. By contrast, in both *L. corniculatus* accessions, Cl^−^ and Na^+^ increased in the stress treatments where irrigation was supplemented with NaCl, both in leaves and roots (Figure 4A-I). Nevertheless, the ion accumulation in both fractions of leaves (young and old) was always higher than in roots, showing that the ions were transported to shoots when plants were exposed to saline and waterlogging-saline stress.

**Figure 4.**
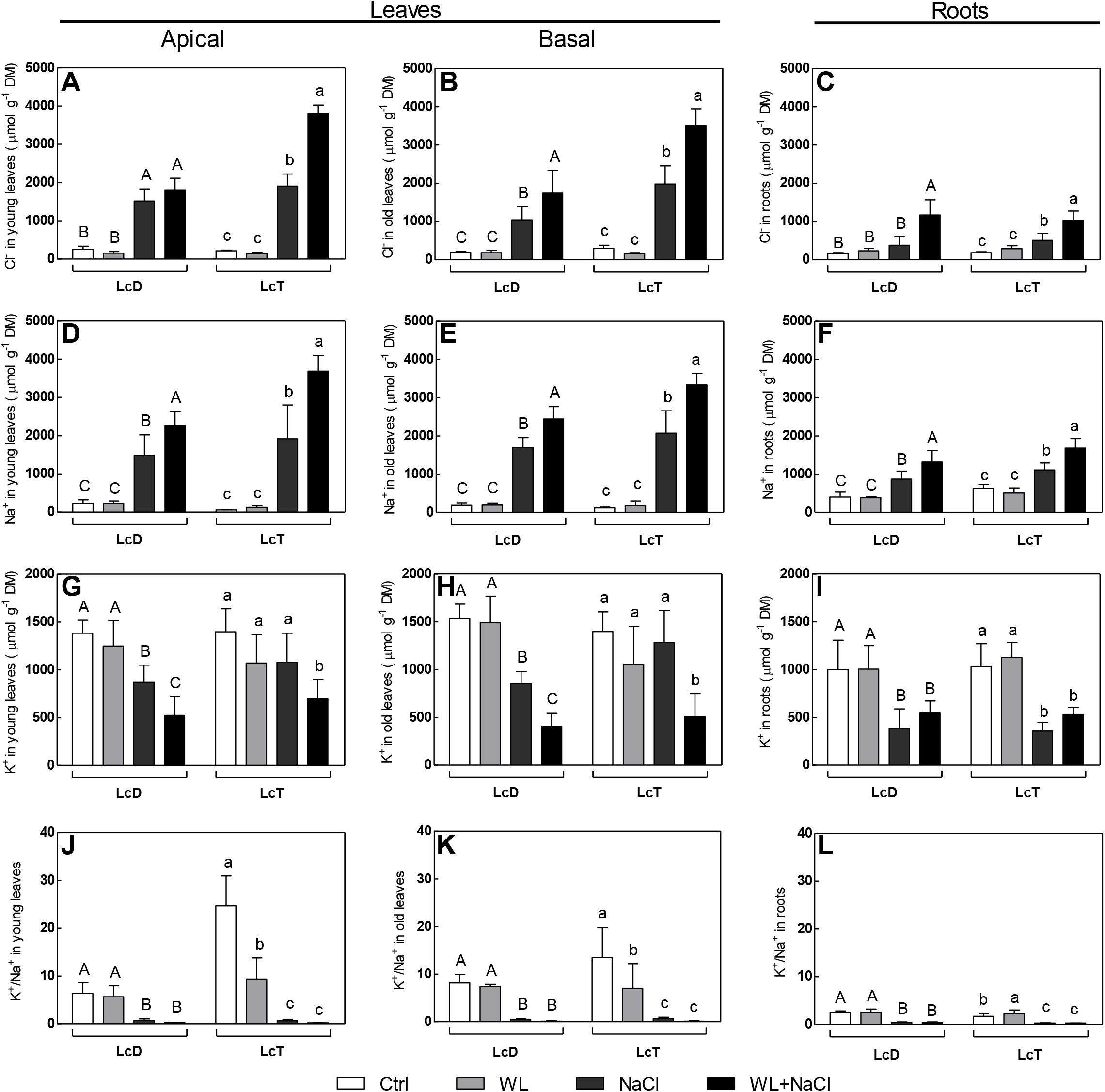
Ions concentrations in apical and basal leaves, and in roots. The concentration of Cl^−^, Na^+^ and K^+^ (μmol.g^−1^.dry mass^-1^) were measured in apical leaves (A, D and G), basal leaves (B, E and H) and in roots (C, F and I) of both *L. corniculatus* accessions under the different treatments (n = 5 ± SD). The K^+^/Na^+^ ratio was calculated for apical and basal leaves (J and K, respectively), and for roots (L), in each case. Means without common letters differ significantly within each of the accessions (one-way ANOVA; Duncan, *p* < 0.05). White bars, control conditions (Ctrl); light gray bars, waterlogging stress (WL); dark gray bars, saline stress (NaCl); black bars, combined stress (WL+NaCl).

For roots and basal leaves tissues, the increase in Cl^−^ and Na^+^ concentration was larger under the combined stress condition, when compared to salinity, in both *L. corniculatus* accessions. Interestingly, Cl^−^ accumulation in apical leaves of LcD was similar between saline and waterlogging-saline stress (Figure 4A). Nevertheless, in general, a similar pattern of ion accumulation was observed between apical and basal leaves of both LcT and LcD; although Cl^−^ and Na^+^ accumulation was higher for LcT, when compared to LcD, under the combined stress treatment (t-test, *p* < 0.05).

Regarding the K^+^ concentration, a reduction in apical and basal leaves was observed in both saline and combined stress treatments, when compared with controls, for LcD. Meanwhile, a significant decreased was only measured under the waterlogging-saline treatment for LcT (Figure 4G and H). In the case of roots, salinity and waterlogging-saline stress reduced the concentration of K^+^ in similar proportions for LcD and LcT, compared with their respective controls (Figure 4I). If we consider the changes in both K^+^ and Na^+^ concentrations, a decrease in K^+^/Na^+^ ratio was also observed under the salt and combined stress conditions for both LcT and LcD, compared to controls (Figure 4J-L). No differences were observed for the K^+^/Na^+^ ratio between the waterlogging stress and control treatments for LcD, neither in leaves nor roots, but differences were detected for LcT.

### 3.4 Effect of stress treatments on the expression of *CLC* and *NHX1* genes

*CLC* and *NHX1* are genes coding for Cl^−^ and Na^+^ transporters, respectively, that have been reported to participate in ion homeostasis and salt stress tolerance in different plant species (Bao et al., 2014; Diédhiou and Golldack, 2006; Jossier et al., 2010; Nakamura et al., 2006; Teakle and Tyerman, 2010). The relative expression of both genes was measured on the most contrasting treatments (control and combined waterlogging-saline stress) for both LcD and LcT (Figure 5). The relative expression of *CLC* and *NHX1* was not affected by waterlogging-salt treatment in LcT plants (Figure 5A and B, respectively). Nevertheless, in the LcD accession, the CLC gene expression increased three-folds under the combined stress condition, when compared to its control (Figure 5A). Regarding the *NHX1* expression, an increased expression trend was observed for LcD, although in this case the difference was not significant (Figure 5B).

**Figure 5.**
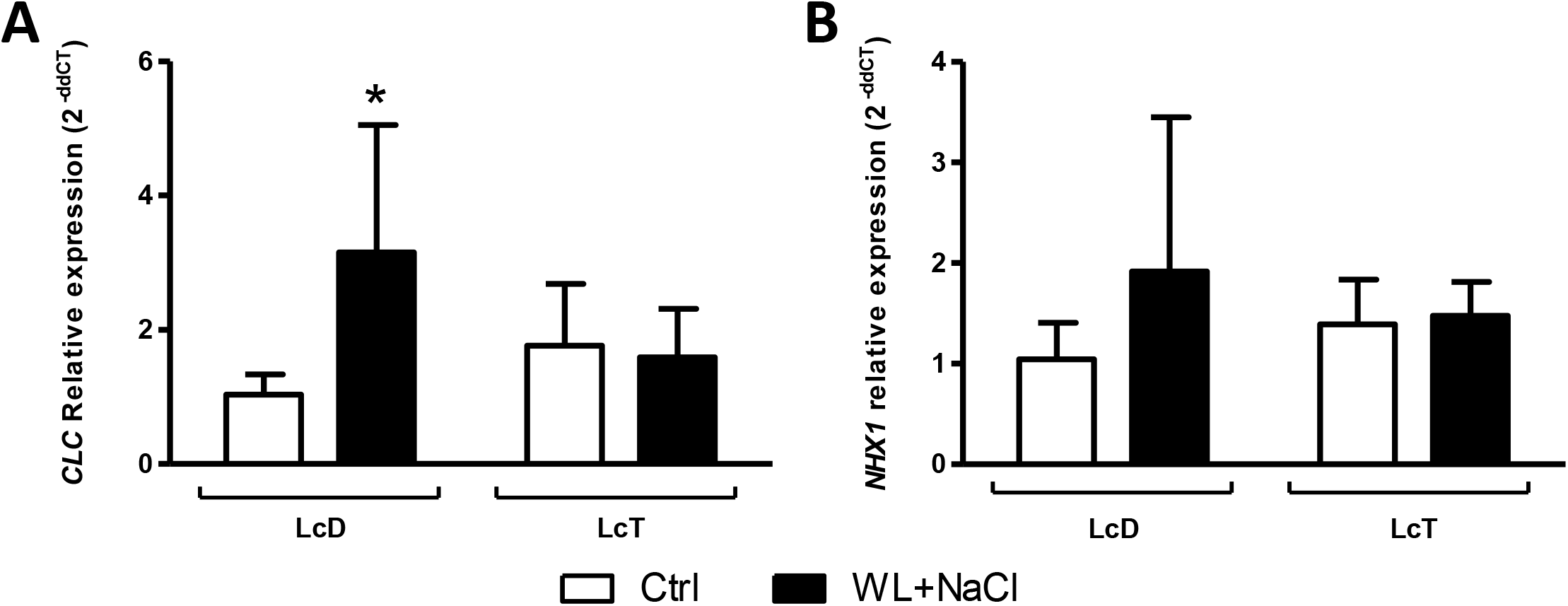
Relative expression of *CLC* (A) and *NHX1* (B) genes. The values represent the mean ± SD (n = 3). Statistical analysis of relative expression was performed by comparing the relative expression of the genes based on the pairwise fixed reallocation randomization test (p < 0.05). Asterisk show significant differences (p < 0.05). White bars, control treatment (Ctrl); Black bars, combined stress (WL+NaCl).

## 4. Discussion

In the present study, the stress tolerance of a recent described *L. corniculatus* diploid accession was addressed, with the hypothesis that, due to the characteristics of its ecological niche (Escaray et al., 2014), it would present a better performance under waterlogging-saline conditions than other *L. corniculatus* accession. The larger variability for LcD can be justified with the fact that it was collected from a natural population and did not go through a domestication process such as the commercial cultivar LcT (Escaray et al., 2014). It is worth mentioning that a high variability within individuals of the same population could be an advantage in the search of tolerant traits for forage breeding programs. In fact, the inter-specific hybridization of LcD with other species was already demonstrated to be a useful tool for improving forage legumes (Antonelli et al., 2019; Escaray et al., 2019, 2014).

As a general response, waterlogging stress significantly affected shoot and root dry mass accumulation and net photosynthetic rate in LcT, but did not cause a severe impairment in growth in neither of both accessions. The tolerance extent of *L. corniculatus* to waterlogging stress was previously described (Antonelli et al., 2019; Striker and Colmer, 2017), and is related with the ability of aerenchyma and adventitious root formation. These traits allow O_2_ supply and respiration, sustaining metabolism and growth rates under the hypoxic conditions imposed by the water submergence (Colmer and Voesenek, 2009; McDonald et al., 2002). Another common plant response to waterlogging is the higher relative partition of photosynthates to shoots than to roots, resulting in an increased shoot:root ratio (Mendoza et al., 2005; Rubio et al., 1995). This was observed for both LcT and LcD under the waterlogging stress treatment.

On the contrary, plants were severely affected under the saline stress treatments. This effect was more evident in the tetraploid accession, showing a decrease in its dry biomass accumulation and in the photosynthetic parameters evaluated. Both measurements are consistent, showing that a reduction in net photosynthetic rate has a significant impact in dry biomass accumulation. Salinity has been described as a two-phase stress, where a first osmotic component affects plant growth, followed by a toxic phase due to the accumulation of toxic ions (Munns, 2002). Due to the salinity acclimation period that was imposed before the salinity treatments (see Materials and Methods), the osmotic phase was probably reduced and the decrease in the photosynthetic capacity of the plants could be attributed to the accumulation of toxic levels of ions. Nevertheless, although an increase in the concentration of Cl^−^ and Na^+^ was observed under salt stress in LcD and LcT, the maximum quantum yield of PSII was not severely affected under this condition. Similar results were reported in the halophyte *Suaeda salsa* (Lu et al., 2002), and for LcD under comparable saline stress conditions (Escaray et al., 2019).

Considering the decrease in stomatal conductivity, an osmotic effect due to salinity cannot be discarded, and the reduction in the photosynthetic capacity of both accessions could be also attributed to a lower CO_2_ availability. Similar results were observed in other species, where it was concluded that stomatal aperture was the main factor limiting leaf photosynthetic capacity in NaCl-treated plants (Loreto et al., 2003; Meloni et al., 2003). The lack of alterations in PSII functionality under salt stress might be the result of an increase in photoprotection mechanisms, such as Non-Photochemical Quenching and cyclic electron flow (Bencke-Malato et al., 2014; Dionisio-Sese and Tobita, 2000), or the increase in the activity of ROS scavenging mechanisms in the chloroplast (Badawi et al., 2004; Meloni et al., 2003). The latter was previously reported to take place in other *Lotus* species, although under low temperature conditions (Calzadilla et al., 2016).

Despite salinity similarly affected both *L. corniculatus* accessions, the larger contrasting response between accessions was observed under the combined stress treatment. This condition mainly affected LcT plants; while for LcD, unexpectedly, waterlogging seemed to reduce the negative effects caused by salinity (Figure 1). The better performance of LcD compared to LcT could be, at least partially, explained by a lower accumulation of Cl^−^ and Na^+^ ions in their leaves. Combined waterlogging-saline stress cause an increase in these ions concentrations when compared to salinity (Barrett-Lennard, 2003), which was, in general, more pronounced for LcT plants.

The saline stress tolerance has been correlated to the capacity of Cl^−^ exclusion in different legume species (Teakle and Tyerman, 2010), including some of the *Lotus* genus (Sanchez et al., 2010; Teakle et al., 2006). Moreover, the negative correlation between Cl^−^ concentration and stress tolerance has been reported to be even stronger than the one existing for Na^+^, in *Trifolium* (Rogers et al., 1997), *Medicago* (Sibole et al., 2003), *Glycine* (Luo et al., 2005) and even *Lotus* (Teakle et al., 2007, 2006). Xu et al. (1999) defined critical values for Cl^−^ toxicity above 200 μmol.g ^_1^.DM for sensitive crops species, such as rice, wheat, alfalfa and peanut; and above 1400 μmol.g^1^.DM for the most tolerant ones, i.e. sugar beet and tomato. Under the combined stress condition, the values measured for LcT were twice as high as the one observed for LcD, although in both cases Cl^−^ concentration reached the levels defined as toxic (Xu et al., 1999). This toxic effect was clearly observed in the reduction of the maximum quantum yield of PSII in LcT, in the further reduction of dry biomass accumulation and in the appearance of chlorotic and senesced leaves in this accession.

By contrast, no symptoms of ion toxicity were observed in LcD under the waterlogging-saline treatment, despite the high levels of Cl^−^ concentration in leaves. Even more, surprisingly, a net photosynthetic rate increased was observed for this accession under the combined stress, when compared with the salinity treatment alone. Although it is known that waterlogging deepens the effects caused by salinity stress (Barrett-Lennard, 2003; Bennett et al., 2009), similar results regarding the amelioration of salinity were recently reported in *Mentha aquatic* (Haddadi et al., 2016). These results were suggested to be the consequence of the priming of an antioxidant response, which could help to increase membrane stability and reduce the toxic effects of NaCl (Haddadi et al., 2016). However, further studies are needed to understand how photosynthesis acclimates to the combined stress condition.

The better tolerance of LcD to the combined stress condition could be explained through a better compartmentalization of toxic ions within the cells. Different subcellular compartmentalization mechanisms have been described in the plant response to saline stress (revised by Munns and Tester, 2008; Teakle and Tyerman, 2010), including the participation of ion transporters such as *CLC* and *NHX1*, for Cl^−^ and Na^+^ respectively (Bao et al., 2014; Jossier et al., 2010; Nakamura et al., 2006). In this sense, expression of *CLC* was strongly correlated with salinity tolerance in grapevine (Henderson et al., 2014), while overexpression of *NHX1* was demonstrated to enhance salt stress in Arabidopsis and even *L. corniculatus* (Liu et al., 2010; Sun et al., 2006). These transgenic plants showed a higher Na^+^ accumulation in their tissues, but a higher photosynthetic capacity. The amelioration of the toxic effect of Na^+^ was ascribed to its compartmentalization into the vacuole (Liu et al., 2010), which provides an efficient way to alleviate Na^+^ excess in the cytosol, and helps keeping cellular turgence under stress (Flowers et al., 1977). Furthermore, overexpression of tonoplast related proteins was also recently associated with a higher tolerance to salinity due to ion compartmentalization in legumes, such as *L. corniculatus* (Bao et al., 2014) and *M. sativa* (Bao et al., 2016).

The gene expression of *CLC* and *NHX1* was measured in order to approach a possible mechanism of waterlogging-saline tolerance in both *L. corniculatus* accessions. The expression of *CLC* was significantly increased under stress in LcD, while a similar expression trend was also observed for *NHX1;* meanwhile, plants of LcT showed no effect in relative expression of both genes between treatments. Chloride channels are involved into intracellular compartmentalization of Cl^−^, sequestering this anion to prevent toxic levels in cytoplasm. Our results suggest that, in LcD, increasing the expression of *CLC* might favour the compartmentalization of Cl^−^, improving its waterlogging-saline stress response. A similar effect might be taking place for Na^+^ and its transporter NHX1. In this sense, a previous work showed that *NHX1* is also involved in *L. tenuis* response to waterlogging-salinity stress, and that its expression levels can justify the better tolerance of *L. tenuis* when compared to *L. corniculatus* (commercial cv. San Gabriel) (Teakle et al., 2010). Nevertheless, in this case, the differences in *NHX1* expression between *L. tenuis* and *L. corniculatus* were shown in root tissues.

It is worth mentioning that the active transport of ions against their concentration gradient implies the consumption of energy (Colmer and Flowers, 2008; Munns and Tester, 2008). Thus, the hypoxic condition imposed by flooding severely affects respiration, the production of ATP and, as a consequence, affects ions compartmentalization as a salinity stress response (Barrett-Lennard, 2003; Kotula et al., 2015). This is one of the reasons why a more severe effect of salinity is generally observed when is combined with waterlogging (Barrett-Lennard, 2003; Teakle et al., 2007). Wetland halophytes plants were reported to have a lower increase in Na^+^ and Cl^−^ ions due to a higher oxygenation of their roots (Colmer and Flowers, 2008). Interestingly, Antonelli et al. (2019) measured the response of different *Lotus* species to waterlogging stress, including the ones addressed in the present study, and found that LcD shows almost three times the percentage of root aerenchyma than LcT. These results could imply a better root oxygenation in the first mentioned accession, which would allow respiration and ATP generation under hypoxic conditions. The higher root aerenchyma formation of LcD, when compared to LcT, could also justify its better response to the waterlogging-saline stress condition. Similar results were obtained for *L. tenuis*, when compared to LcT, by Teakle et al. (2007).

There is a general agreement that cytoplasmic high K^+^/Na^+^ ratio is a good indicator of low salt damage and high salinity tolerance (Maathuis and Amtmann, 1999; Munns and Tester, 2008). The similar physicochemical characteristics between K^+^ and Na^+^ affect a wide range of metabolic processes, such as enzymatic reactions and protein synthesis, and maintaining the K^+^/Na^+^ ratio under stress conditions is of key importance to maintain K^+^ homeostasis (Almeida et al., 2017; Maathuis and Amtmann, 1999; Shabala and Cuin, 2008). However, it was previously reported that for halophytes and some glycophytes plants, the K^+^/Na^+^ ratio is not a good parameter to assess salinity tolerance (Colmer and Voesenek, 2009).

Our results show that, in leaves, the K^+^/Na^+^ was only reduced in LcD when the irrigation solution was supplemented with NaCl; while in LcT, K^+^/Na^+^ was decreased in all of the stress treatments imposed. These results are in agreement with a better response of LcD to waterlogging, when compared to LcT (our own results; Antonelli et al., 2019). Nevertheless, for the salinity and combined stress treatments, the K7Na^+^ ratio values did not differ between the evaluated *Lotus* accessions. Thus, the K^+^/Na^+^, at least at tissue level, is not a strong indicator of salinity tolerance for species of the *Lotus* genus, in agreement with results obtained by other authors (Rejili et al., 2007).

Although salinity stress reduces K^+^ and increases Na^+^ concentration in plant tissues, this not necessarily implies changes in their cytoplasmic concentrations (Flowers et al., 2015). For instance, a significant proportion of K^+^ of the leaves is located in the vacuole and is responsible of keeping cell turgence (Andrés et al., 2014; Barragán et al., 2012). In certain plant species, such as *Mesembryanthemum cristallinum* and *Suaeda maritima* (and probably *Lotus*), the role of K^+^ could be replaced by Na^+^, which would allow maintaining ion homeostasis in the cytosol (Kronzucker et al., 2013; Leigh and Wyn Jones, 1986). As a consequence, for plants with a high salinity response implying subcellular ion compartmentalization, changes in the K^+^/Na^+^ ratio might not necessarily imply alteration in K^+^ homeostasis and stress sensitivity.

## 5. Conclusions

In the present study, the stress tolerance to waterlogging, salinity and combined waterlogging-saline stress was evaluated in two accessions of *Lotus corniculatus*. Our results showed that the diploid accession, obtained from an environmental niche naturally affected by waterlogging and salinity, has a better response to all the stress conditions evaluated, when compared to LcT. This contrasting response was more evident under the combined stress treatment. A lower decrease in dry biomass accumulation and absence of stress symptoms were observed in treated LcD plants, when compared with LcT, which could be ascribe to a lower photoinhibitory effect and lower Cl^−^ and Na^+^ accumulation in leaves. In addition, the better response could be justified trough the triggering of ion subcellular compartmentalization mechanisms, which was suggested through the increased expression levels of the *CLC* and *NHX1* transporters genes. In this sense, we suggest that the K^+^/Na^+^ ratio is not a good indicator of salinity tolerance in plants where ion compartmentalization responses take place. As a conclusion, the higher adaptability of the *L. corniculatus* diploid accession to combined waterlogging-saline stress was demonstrated when compared to another *L. corniculatus* commercial cultivar. Thus, the recently characterized *L. corniculatus* accession could be used to introduce new tolerant traits to waterlogging-saline stress, in other *Lotus* species commonly used as forage.

## Acknowledgements

This work was supported by grants from the *Agencia Nacional de Promoción Científica y Tecnológica* (ANPCYT-Argentina)/FONCyT-PICTs 1560, 1612, 3648 and 3718 and *Consejo Nacional de Investigaciones Científicas y Técnicas* (CONICET-Argentina)/ PIP 0980. C.J.A. and P.I.C. and M.P.C were doctoral fellows, whereas F.J.E. and O.A.R. are members of the Research Career of CONICET.

## Authors contribution

C.J.A. designed and performed all the experiments, and analyzed data; P.I.C. analyzed data; M.P.C performed some experiments and analyzed data; F.J.E design experiments and analyzed data; O.A.R. conceived the project, designed and supervised all the experiments. The article was written by O.A.R, C.J.A. and P.I.C.

## Supplementary material

**Supplementary Figure 1.**
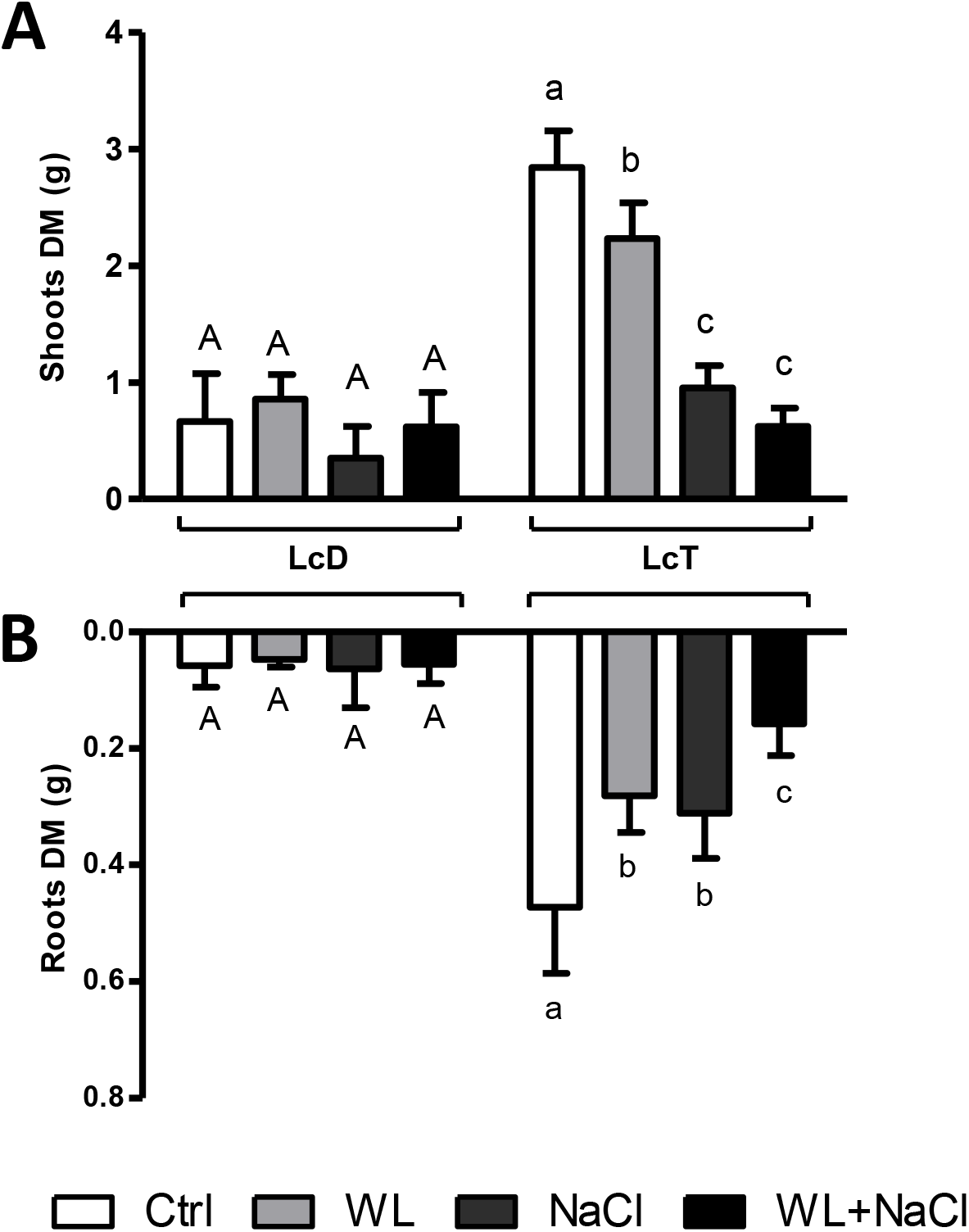
Shoots and roots dry mass accumulation of L. corniculatus accessions subjected to the different stress treatments. The measured shoots (A) and roots (B) dry mass is shown in grams (n = 5 ± SE). Means without common letters differ significantly within each of the accessions (one-way ANOVA; Duncan, p < 0.05). White bars, control conditions (Ctrl); light gray bars waterlogging stress (WL); dark gray bars, saline stress (NaCl); black bars, combined stress (WL+NaCl).

**Supplementary Figure 2.**
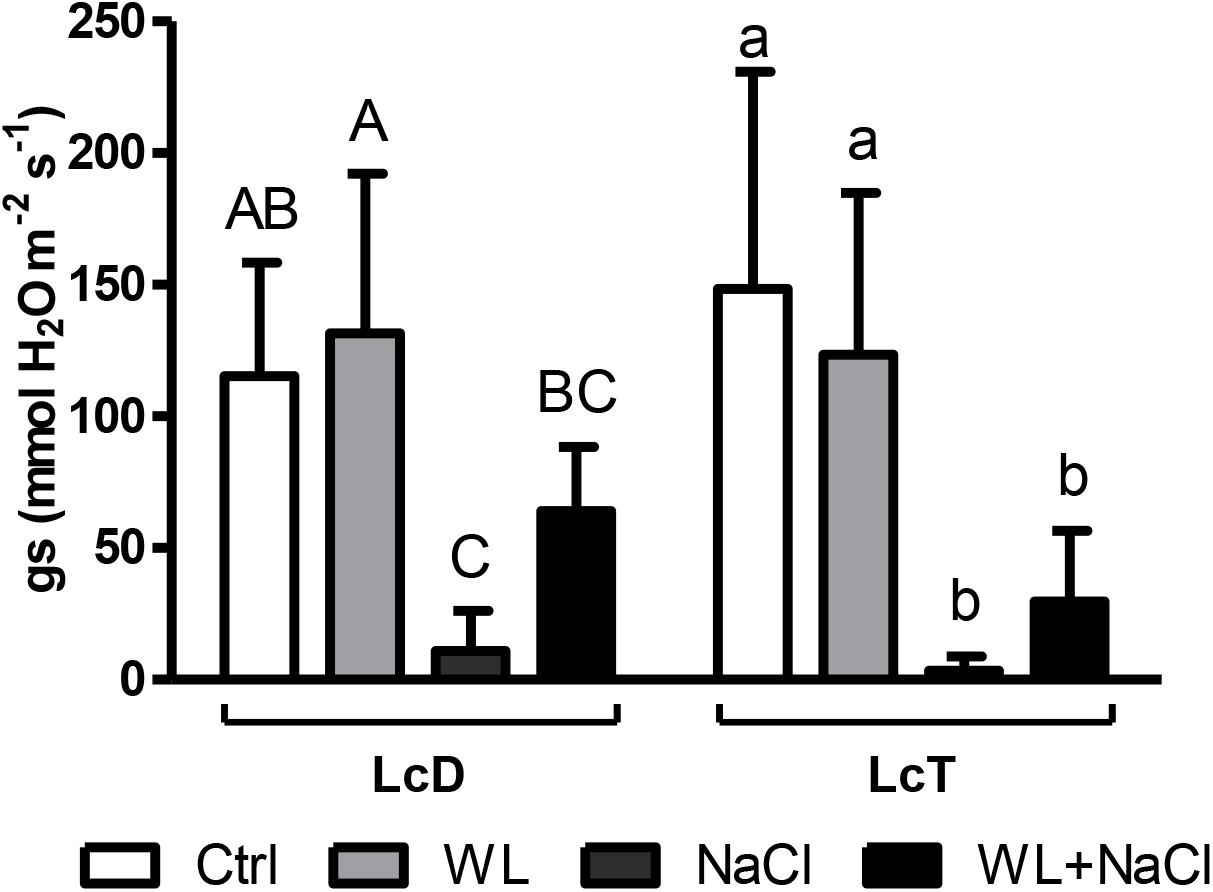
Stomatal conductance of the L. corniculatus accessions under the different stress treatments. The stomatal conductance was measured for five independent biological replicates (n = 5 ± SD). Means without common letters differ significantly within each of the accessions (one-way ANOVA; Duncan, p < 0.05). White bars, control conditions (Ctrl); light gray bars waterlogging stress (WL); dark gray bars, saline stress (NaCl); black bars, combined stress (WL+NaCl).

**Supplementary Table 1.**
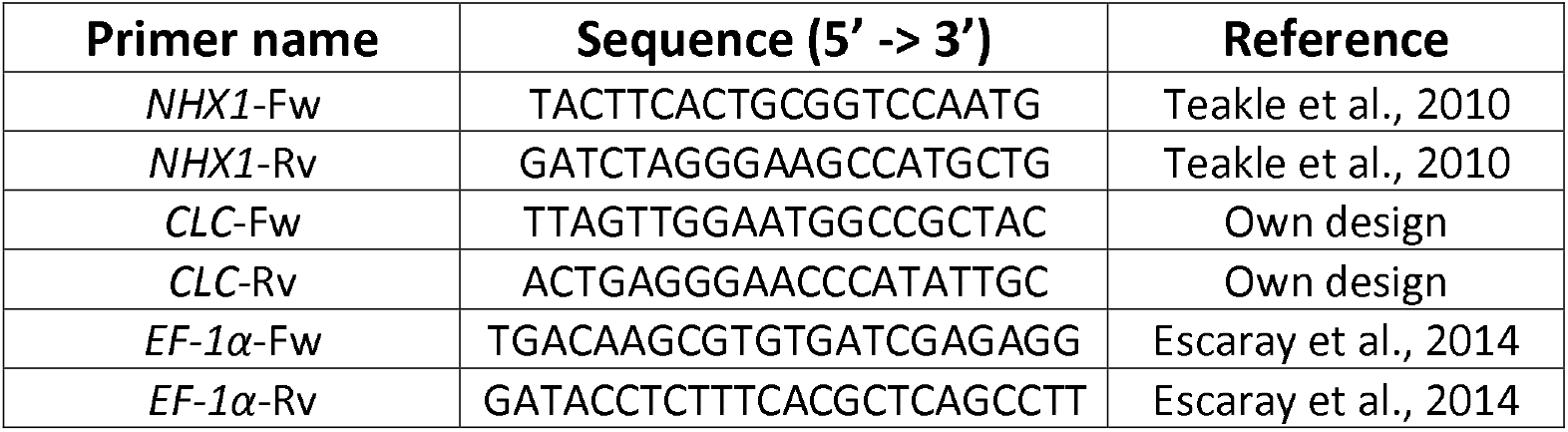
Primers used for qRT-PCR.

